# Adaptive siderophore repression and redox-metabolic remodeling limit *E. coli* Fenton-mediated cytotoxicity after H_2_O_2_ exposure

**DOI:** 10.64898/2026.09.17.752378

**Authors:** Elise Bulaoro, Bharat Mishra, Anuradha Goswami

## Abstract

Hydrogen peroxide imposes linked redox and metal stress on bacteria by promoting ferrous iron-dependent Fenton chemistry, generating reactive species that damage DNA, proteins, iron-sulfur cofactors, and membranes. Because siderophores expand iron acquisition, their continued production during peroxide exposure may increase cytotoxicity by enlarging the labile iron pool. Here, we identify an adaptive iron-restriction response that connects siderophore repression, envelope integrity, and metabolic remodeling during bacterial peroxide stress. Using *Acinetobacter baumannii*, *Escherichia coli*, *Pseudomonas aeruginosa*, and *Staphylococcus aureus*, we evaluated cellular oxidation kinetics, growth-normalized extracellular siderophore release, membrane permeabilization, and viability following sub-inhibitory H_2_O_2_ exposure. Acute peroxide treatment significantly elevated general fluorophore oxidation across all species (p < 0.0001), while prolonged exposure reduced biomass-normalized siderophore output, most strongly in *E. coli* and *S. aureus*. Membrane permeability accounted for a substantial fraction of extracellular siderophore variation, indicating that envelope alterations link peroxide-induced damage to iron-acquisition dynamics. We further integrated these physical phenotypes with differential transcriptomics, oxidation gene expression regression, E-Flux-constrained genome-scale metabolic modeling, flux variability analysis, shadow price analysis, and in silico gene-essentiality mapping in acute and 500-generation H_2_O_2_-adapted *E. coli*. Acute stress transiently co-activated iron uptake, efflux, NADPH regeneration, and stress-defense genes. In contrast, long-term adaptation systematically repressed enterobactin biosynthesis and uptake components (*fepA*, *entE*, *fes, ryhB*) while inducing persistent peroxide-and acid-resistance modules (*ahpC*, *gadA*, *gadB*). Genome-scale metabolic modeling independently verified that this evolutionary adaptation contracted flux variability, relieved envelope bottlenecks, and eliminated acute stress-specific single-gene vulnerabilities at the cost of reduced maximal growth capacity. Together, these results define siderophore repression as a system-level adaptive response that limits Fenton-reactive iron influx while reallocating metabolic resources toward redox buffering, membrane stabilization, and genetic robustness.

## 1. Introduction

Fenton chemistry has progressed from a nineteenth-century synthetic chemistry discovery to a foundational concept in redox biology ^1^. In 1894, Henry John Horstman Fenton showed that hydrogen peroxide (H_2_O_2_) reacts with ferrous iron (Fe^2+^) to produce a strongly oxidizing reaction ^2^. Later mechanistic work by Haber, Weiss, and colleagues demonstrated that this reaction generates highly reactive, non-selective hydroxyl radicals (^•^OH): Fe^2+^ + H_2_O_2_ → Fe^3+^ + OH^—^ +• OH ^3^. The biological significance of this reaction became evident after McCord and Fridovich discovered superoxide dismutase, establishing that aerobic cells continuously produce reactive oxygen species (ROS), including superoxide (O_2_ ^•—^) and H_2_O_2_, as metabolic by-products ^4^. This insight connected inorganic radical chemistry to intracellular oxidative stress and identified iron homeostasis as a key determinant of bacterial survival.

Hydrogen peroxide is generated naturally during metabolism and can form highly reactive hydroxyl radicals (^•^OH)[Reactive Oxygen Species (ROS)], particularly when it reacts with intracellular labile Fe^2+^ through the Fenton reaction. Although bacteria use catalases, peroxidases, superoxide dismutases, ferritins, and related defense systems to control physiological ROS levels ^5–7^, widespread use of H_2_O_2_ in healthcare ^8^, food sanitation ^9^, and wastewater treatment ^10^ can expose environmental and clinical isolates to sub-lethal oxidative stress ^10,11^. In bacteria, this stress can damage DNA, iron-sulfur (Fe-S) clusters, enzymes, and membrane lipids ^5^. To prevent catastrophic Fenton-mediated hydroxyl radical (•OH) surges, bacteria maintain a delicate balance between iron acquisition, storage, and ROS scavenging ^12^. Although siderophore secretion is classically defined as a high-affinity iron acquisition strategy, recent hypotheses have debated whether active chelator release contributes to antioxidant redox buffering or acts as pro-oxidant liability under oxidative stress^13–15^ . Iron imports directly expand the intracellular labile iron pool, continued siderophore synthesis under peroxide challenge poses a severe threat of Fenton-driven lethality. Therefore, we hypothesize that iron acquisition during peroxide (H_2_O_2_) exposure results in cytotoxicity due to Fenton chemistry, and thus bacteria adapt to repress or silence siderophore production under long-term oxidative stress exposure.

This study investigated how peroxide (H_2_O_2_) stress drives bacteria to downregulate siderophore production and undergo redox perturbation as a defensive metabolic adaptative strategy. We define the dynamic coupling among iron homeostasis, oxidation increase, and metabolic adaptation across four major bacterial pathogens (*Acinetobacter baumannii*, *Escherichia coli*, *Pseudomonas aeruginosa*, and *Staphylococcus aureus*) exposed to sub-minimal inhibitory concentrations (sub-MICs) of H_2_O_2_. Specifically, we examine how iron-mediated regulation, cross-protection, and central metabolic shifts support bacterial survival during prolonged peroxide stress. Using phenotypic assays that track viability, extracellular siderophore release, real-time intracellular oxidation kinetics, and membrane permeability, we characterize species-specific physiological adaptations to oxidative stress. To define the molecular mechanisms underlying these adaptive phenotypes, we integrated multi-species experimental profiling with transcriptomic and flux balance analyses utilizing publicly available RNA-seq datasets for *Escherichia coli* MG1655 from Zorraquino et al., 2017 ^16^. The regulatory architecture was characterized across two distinct oxidative regimes: *(i)* acute unevolved shock, comparing baseline wild-type cells against those abruptly exposed to H_2_O_2_ to capture immediate transcriptional and metabolic rewiring; and *(ii)* long-term evolutionary adaptation, comparing acute exposure against an H_2_O_2_ adapted strain to resolve how sustained selective pressure reshapes metabolic flux, optimizes NADPH supply-demand balance, and minimizes endogenous ROS generation. By coupling phenotypic trajectories across diverse bacterial species with transcriptome and metabolic modeling in *E. coli*, this study uncovers oxidation–siderophore interactions and global metabolic pivots, demonstrating a coordinated axis between iron homeostasis and central metabolism that promotes bacterial survival and cross-protection under oxidative challenge.

## 2. Method

### 2.1. Bacterial Strains and Growth Conditions

*Acinetobacter baumannii* ATCC 19606, *Escherichia coli* MG1655, *Pseudomonas aeruginosa* PAO1, and *Staphylococcus aureus* USA300 were cultured aerobically in Luria-Bertani (LB) broth at 37°C under batch conditions. Cultures were exposed to species-specific sub-minimal inhibitory concentrations (sub-MICs) of hydrogen peroxide (H_2_O_2_) for either 1 h, representing acute oxidative shock, or 24 h, representing sustained oxidative-stress exposure. Because H_2_O_2_ can degrade during prolonged batch incubation, the 24 h time point was interpreted as the downstream physiological state established after sustained peroxide challenge rather than continuous exposure to a constant oxidant concentration. Untreated control cultures were maintained under identical conditions without H_2_O_2_. The sub-MIC value of each isolate was determined after growing bacteria overnight (starting culture OD_600_ adjusted to 0.06) in increasing concentration of H_2_O_2_ spiked LB broth till no visible growth was observed; the highest concentration permitting visible growth was defined as sub-MIC for this study.

### 2.2. Oxidative Stress Batch Experiments

Fresh overnight cultures of *A. baumannii*, *E. coli*, *P. aeruginosa*, and *S. aureus* were diluted 1:100 into sterile LB broth and grown in 50 mL sterile flasks at 37°C with shaking at 225 rpm for 3 h to obtain mid-logarithmic-phase cultures. For each strain, 15 mL cultures were assigned to treatment groups in biological triplicate (n = 3). Untreated growth controls and media-only negative controls were included in parallel. Oxidative stress was induced by adding H_2_O_2_ to the following final sub-MIC concentrations: *A. baumannii*, 22.93 mM; *E. coli*, 8.53 mM; *P. aeruginosa*, 35.28 mM; and *S. aureus*, 276.36 mM. The higher concentration used for *S. aureus* reflected its greater intrinsic tolerance to acute peroxide stress relative to the other isolates tested. Samples were collected immediately before treatment (T = 0), after 1 h of exposure (T = 1 h), and after 24 h of incubation (T = 24 h). At each time-points, 5 mL aliquots were collected from treated and control cultures, bacterial growth was monitored by measuring OD_600_ using a microplate reader, and aliquots were processed immediately by washing in sterile buffer to remove remaining H_2_O_2_ in media and process cell pellet for the downstream phenotypic assays.

#### Viable Cell Counts and Antimicrobial Susceptibility Testing

Bacterial viability was quantified using standard plate-count enumeration. Pellets obtained after three washes were serially diluted, and 10 µL aliquots of selected dilutions were plated on appropriate agar media and incubated at the optimal growth temperature. Colony-forming units were counted and expressed as CFU/mL using the equation: CFU/mL = number of colonies × dilution factor / volume plated. Viable colonies recovered from each experiment were subsequently used for antimicrobial susceptibility testing by the Kirby-Bauer disk diffusion method ^17^. Briefly, 10 µL of overnight LB culture was spread uniformly across Mueller-Hinton agar plates using a sterile cotton swab to generate a bacterial lawn. Antibiotic-impregnated disks were placed on the inoculated agar surface, and plates were inverted and incubated at 37°C for 24 h. Susceptibility was assessed by measuring the diameter of the zone of inhibition around each disk and comparing values with Clinical and Laboratory Standards Institute (CLSI) ^18^.

#### Intracellular Oxidative Quantification

Intracellular oxidation was measured through fluorescence using the Invitrogen Thermo Fisher Scientific Reactive Oxygen Species Fluorometric Assay Kit (Green). Bacterial cells were incubated with 2′,7′-dichlorodihydrofluorescein diacetate (DCFH-DA) during logarithmic growth to allow dye incorporation ^19^. Excess extracellular dye was removed by washing, and labeled cells were resuspended in LB medium containing the appropriate sub-MIC concentration of H_2_O_2_. Cells were incubated in the dark according to the manufacturer’s protocol. After washing, fluorescence was measured as relative fluorescence units (RFU) using a microplate reader with excitation at 488 nm and emission at 525 nm. While general fluorometric probes (e.g., DCFH-DA) monitor cellular oxidation kinetics rather than specific ROS species, we relied on an integrated phenotype-to-model approach—combining membrane permeability, global transcriptomic profiling, and genome-scale metabolic flux modeling—to define cellular oxidative stress responses independently of probe-specific kinetics.

#### Siderophore Detection

Extracellular siderophore production was measured using the universal Chrome Azurol S (CAS) colorimetric assay^20^. The CAS reagent forms a blue complex with ferric iron and hexadecyltrimethylammonium bromide. Culture supernatants collected after treatment were mixed with CAS reagent; siderophore-mediated iron removal from the CAS complex produced a color shift from blue to orange/yellow. Siderophore activity was quantified by measuring absorbance at 630 nm, where lower absorbance indicated greater siderophore-mediated iron chelation. Percent siderophore units (%PSU) were calculated as: %PSU = [(A_control_ − A_sample_) / A_control_] × 100, where A_control_ represents the absorbance of the negative control (CAS + LB) and A_sample_ represents the absorbance of the CAS-supernatant mixture.

#### Cell permeability assay using Propidium Iodide

Bacterial cells were stained with Propidium Iodide (PI) to measure the cellular oxidative damage^21^. Aliquot of 100 μL of washed pellet suspension before and after H_2_O_2_ experiment was added to black-walled 96-well plate. 3 µL of the Propidium Iodide (PI) Stock Solution (1.0 mg/mL) was directly to each 100 µL cell sample. Unstained cells (negative control) and cells treated with a permeabilizing agent Triton X-100 were used as positive control. Plates were incubated in dark at room temperature for 10 min and fluorescence was recorded in Relative Fluorescence Unit (RFU) using microplate reader at excitation wavelength 535 nm and emission wavelength at 617 nm.

### 2.3. Escherichia coli transcriptomic processing and differential expression analysis

Publicly available RNA-seq data from Zorraquino et al., 2017 ^16^ were obtained from NCBI GEO accession GSE58325 to characterize genome-wide transcriptional reorganization associated with oxidative-stress resilience in *E. coli*. In the original study, *E. coli* MG1655 was cultured in M9 minimal salt medium supplemented with 0.4% (w/v) glucose and exposed to 100 mM H_2_O_2_. In this study, two non-overlapping comparisons were analyzed to distinguish acute and evolved responses: (i) an acute unevolved stress-response comparison between baseline wild-type control samples cultured in M9 glucose medium (SRX581894, SRX581901) and wild-type cells abruptly shifted to 100 mM H_2_O_2_ (SRX581911, SRX581918); and (ii) a long-term adaptive comparison using evolved H_2_O_2_-adapted samples (SRX581930, SRX581931). The acute comparison was used to identify immediate transcriptional rewiring during peroxide shock, whereas the evolved comparison was used to evaluate how long-term adaptation reconfigured metabolic pathways, energy allocation, and NADPH demand.

Secondary analysis of bulk RNA-seq FASTQ files was performed using the nf-core/rnaseq pipeline^22^ with the latest available reference genome. The workflow included initial quality control, read trimming, alignment, quantification, and post-alignment quality assessment. Tertiary analysis included additional quality control, normalization, and differential gene-expression analysis. Raw gene counts from acute pre-evolution and post-evolution transcriptomic datasets were normalized to counts per million (CPM) to account for differences in library size across the control, acute-shock, and adapted-state conditions. Log fold changes for acute shock (*LFC_Acute_*) and the adapted state (*LFC_Adapted_*) were calculated. Differential gene expressions were assessed using independent two-sample Welch’s t-tests, with significance defined as *p* < 0.05 and |LFC| ≥ 1.0.

### 2.4. Integrated transcriptomic and Oxidation kinetics modeling

The relationship between intracellular oxidation increases and transcriptional regulation was assessed by integrating DCFH-DA fluorescence measurements with condition-specific gene-expression profiles. Mean fluorescence trajectories across biological replicates were normalized to a relative scale and used as explanatory predictors in ordinary least-squares (OLS) regression models. These models were used to evaluate whether oxidation kinetics predicted gene-expression patterns across oxidative-stress, siderophore, transport, and central-metabolic functional categories.

### 2.5. Genome-Scale Metabolic Model and Environmental Constraints

Constraint-based metabolic modeling was performed using the *E. coli* genome-scale metabolic reconstruction iML1515 ^23^ implemented in COBRApy (v0.29.0+)^24,25^. Baseline physiological conditions were defined by setting maximum lower-bound uptake rates for D-glucose Exchange to −10.0 mmol·gDW^−1^·h^−1^ and molecular Oxygen Exchange reaction to −20.0 mmol·gDW^−1^·h^−1^. To impose condition-specific transcriptomic constraints on reaction capacities, an extended E-Flux strategy^26^ was applied. Differential-expression datasets containing relative fold-change values for acute shock (G0; Fold_Acute_) and long-term adaptation (G500; Fold_Adapted_), each calculated relative to the baseline wild-type control, were mapped to corresponding model gene identifiers.

### 2.6. Phenotypic Growth Prediction and Flux Variability Analysis (FVA)

Maximum theoretical growth rates (μ_max_) were estimated for each physiological state using parsimonious flux balance analysis (pFBA), which minimizes total flux while optimizing biomass production. To evaluate global metabolic flexibility and pathway streamlining across evolutionary states, flux variability analysis (FVA)^27^ was performed at a 95% suboptimal biomass threshold (f = 0.95). For each reaction *r*, the allowable minimum (v_r,min_) and maximum (v_r,max_) flux values were determined, and the flux span was calculated as Δv_r_ = v_r,max_ − v_r,min_. Pathway narrowing was quantified as Δv_r,reduction_ = Δv_r,Acute_ − Δv_r,Adapted_. Inactive reactions were filtered out by setting reactions thershod with active flux ranges (FVA_Span_ > 10^−6^) to be selected and retained for downstream analysis. In Silico single-gene deletion analyses^28^ were conducted in parallel to identify condition-specific essentiality patterns.

### 2.7. Shadow price analysis for bottleneck identification and in-silico single-gene essentiality screening

Rate-limiting metabolites and thermodynamic bottlenecks were identified using linear-programming dual shadow price analysis during biomass optimization^29,30^. Shadow prices represent the sensitivity of the biomass objective to metabolite availability, with non-zero values indicating metabolites that constrain growth. Metabolites with |λ_i_| > 10^−5^ were classified as system-level bottlenecks. Systemic vulnerabilities were further assessed by performing single-gene deletion simulations across acute and adapted metabolic models. In-silico mutant growth rates (v_del_) were compared with the corresponding unmutated condition growth rate (v_WT_). Genes were classified as essential when deletion reduced predicted growth to <10% of the respective wild-type condition (μ_mutant_ < 0.1 × μ_WT_). Essential genes were then stratified as core essential, if required in both acute and adapted states; acute-specific, if essential only during acute shock; or adapted-specific, if essential only after long-term adaptation.

### 2.8. Computational Environment and Visualizations

All linear-programming optimizations were solved using the default GLPK solver in COBRApy. Data visualization was performed in Python using matplotlib (v3.8+) and seaborn (v0.13+), with figures exported at 300 DPI using vector-compatible formatting where appropriate. Additional plots were generated in R version 4.5.2 and Python v3.8+.

## 3. Results

### 3.1. Extracellular siderophore releases were selectively suppressed in a species-dependent manner under oxidative stress

Bacterial siderophores are low-molecular-weight iron-chelating molecules that mediate iron acquisition. In addition to iron scavenging, accumulating evidence indicates that siderophores can influence bacterial responses to oxidative stress^31^. To determine how different bacterial species respond to acute oxidative stress, intracellular oxidation change was monitored in real time during H_2_O_2_ exposure (Figure 1a(i)). All four isolates tested (*A. baumannii*, *E. coli*, *P. aeruginosa*, and *S. aureus*) showed significantly higher fluorescence than untreated controls (*p* < 0.0001). However, both the magnitude and oxidation rate differed across species and between Gram-positive and Gram-negative bacteria (Figure 1a(ii)). The Gram-positive species *S. aureus* showed the greatest relative increase, with a rapid 2.3-fold rise from control. Among the Gram-negative species, *E. coli* exhibited a sustained 1.78-fold increase and reached high total fluorescence levels (>4.5 log_10_ RFU), whereas *P. aeruginosa* increased rapidly before plateauing at 1.52-fold. In contrast, *A. baumannii* displayed a more modest but steady 1.31-fold increase in oxidation rate.

**Figure 1.**
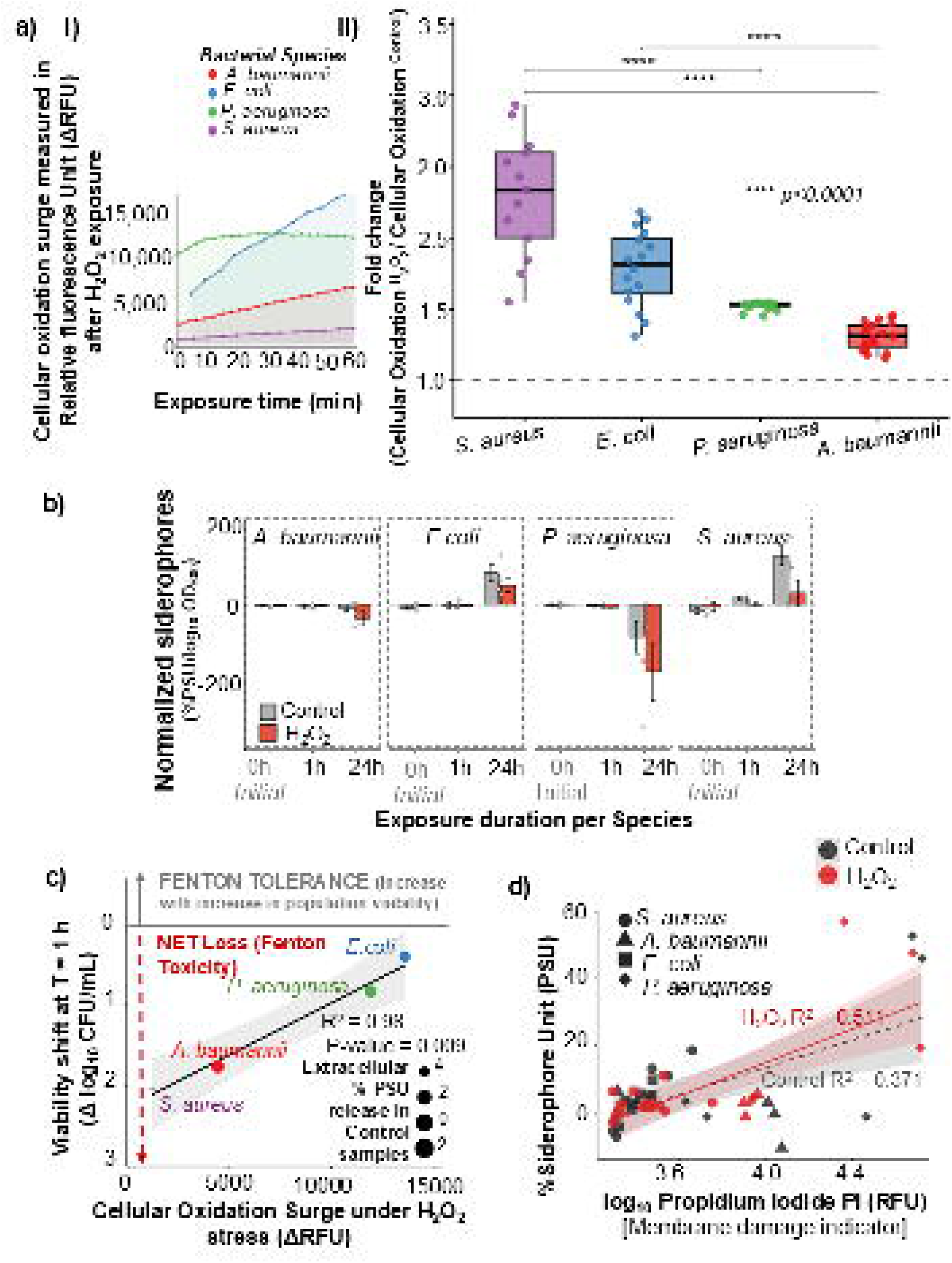
Interspecies cellular oxidation dynamics, siderophore secretion, and Fenton-mediated viability trade-offs under hydrogen peroxide stress. **(a)** Relative intracellular oxidation surge across *S. aureus*, *E. coli*, *P. aeruginosa*, and *A. baumannii*. (i) Kinetic tracking of net oxidation increase, fluorescence measured in Relative Fluorescence Unit (ΔRFU) over a 60 min exposure window. (ii) Fold-change in intracellular Oxidation activity (Cellular Oxidation _H2O2_/Cellular Oxidation _Control_) across species (\*\*\**p* < 0.0001). **(b)** Species-specific normalized siderophore production (%PSU/log_10_OD_600_) under control (gray) and H_2_O_2_-stressed (orange) conditions at baseline (0 h (initial before starting treatment), 1 h, and 24 h. **(c)** Correlation between net cellular oxidation surge (ΔRFU) and population viability shift (Δlog_10_CFU/mL) at T = 1 h (R^2^ = 0.98, p = 0.009). Bubble size corresponds to extracellular percent siderophore unit release in control conditions. **(d)** Relationship between siderophore production (%PSU) and cell membrane disruption (log_10_ Propidium Iodide RFU) under control (gray, R^2^ = 0.371) and H_2_O_2_ treatment (red, R^2^ = 0.511).

Next, to uncover whether oxidative shock (via H_2_O_2_ addition) directly alters iron-acquisition activity independently of biomass, siderophore production was normalized to bacterial growth (log_10_OD_600_) (Figure 1b). Three-way ANOVA identified significant main effects of bacterial species (*p* < 0.001, generalized η^2^ = 0.463) and oxidative treatment (*p* < 0.022, generalized η^2^ = 0.105). Exposure duration alone was not a significant independent predictor (*p* < 0.908); however, significant two-way interactions were observed between species and timepoint (*p* < 0.001, generalized η^2^ = 0.631); oxidative treatment and timepoint (*p* < 0.006, generalized η^2^ = 0.191). Across species, baseline controls and 1 h oxidative-shock conditions showed minimal variation in normalized iron chelator capacity, whereas pronounced species-specific responses emerged after 24 h of exposure (Figure 1b), indicating adaptive response of recovery cells.

Attenuation of extracellular release of siderophores under oxidative stress was especially evident in *E. coli* and *S. aureus*. Unperturbed 24 h *E. coli* cultures showed elevated normalized siderophore secretion (80.42 ± 21.72 % PSU per biomass [log_10_OD_600_]), which decreased by 39.5% under H_2_O_2_ treatment (48.62 ± 16.13 % PSU per biomass [log_10_OD_600_]). *S. aureus* control cultures at 24 h, exhibited strong normalized siderophore activity (123.85 ± 24.42 % PSU per biomass [log_10_OD_600_]). In contrast, H_2_O_2_ exposure reduced normalized siderophore output by 75.7% (30.14 ± 32.39 % PSU per biomass [log_10_OD_600_]).

*P. aeruginosa* exhibited negative capacity ratios at 24 h under both control and peroxide-treated conditions (control: −83.01 ± 41.82; H_2_O_2_: −171.27 ± 74.80 % PSU per biomass [log_10_OD_600_]), suggesting that either biomass accumulation could have exceeded detectable extracellular chelator pools which were further reduced by 24 h incubation. *A. baumannii* showed a similar reduction in normalized capacity during 24 h H_2_O_2_ challenge (control: −8.38 ± 8.38; H_2_O_2_: −36.06 ± 18.12 % PSU per biomass [log_10_OD_600_]). These findings provided us with an in-vitro indication for our hypothesis suggesting that bacteria can putatively attenuate extracellular siderophore release under oxidative stress as a defense strategy to maintain redox homeostasis. Because siderophore-mediated iron import can intensify intracellular Fenton chemistry (Fe^2+^ + H_2_O_2_ → Fe^3+^ + OH^−^ + ^•^OH), reducing siderophore production per unit biomass may function as a protective strategy to limit hydroxyl-radical formation during oxidative stress.

### 3.2. Early viability maintenance and siderophore shutoff are related to intracellular ROS dynamics

Bacterial oxidative stress arises from redox imbalance causes cellular damage^32^. To determine how initial oxidative stress impacts immediate viability and subsequent iron-acquisition behavior, early viability shifts (Δlog_10_CFU/mL) at 1 h post-H_2_O_2_ exposure were correlated against intracellular ROS kinetics and plotted with baseline (control) extracellular siderophore release(Δ %PSU), Figure 1 c. Net intracellular ROS surge predicts early survival capacity where mean internal ROS increase in cells after H_2_O_2_ exposure correlated with siderophore attenuation. Regression analysis revealed a significant linear relationship between intracellular oxidation surge (Δ RFU) at 1 h and short-term viability maintenance during this exposure time (R^2^= 0.98, *p = 0.009*), suggesting redox-dependent survival thresholds. Notably, species displaying low net ROS accumulation, such as *S. aureus* (Δ RFU ≈ 1,200) and *A. baumannii* (Δ RFU ≈ 4,400), suffered the most severe 1 h viability drop, reflecting acute susceptibility to Fenton (Fe^2+^ + H_2_O_2_ → Fe^3+^ + OH^—^ + ^•^OH) mediated cytotoxicity. In *S. aureus*, the lower fluorescent signal (RFU) indicates that the intracellular hydrogen peroxide pool is rapidly depleted and converted by free labile iron into highly destructive hydroxyl radicals (^•^OH). Because ^•^OH are exceptionally reactive and usually exhibit near-diffusion-limited reaction rates with organic molecules (often approaching rate constants of 10^9^ to 10^10^ M^−1^s^−1^)^33^, can cause instantaneous, irreversible macromolecular damage, before they can diffuse or react with fluorescent dyes. Therefore, driving acute lethality despite maintaining a low net fluorescent profile. Conversely, *P. aeruginosa* (Δ RFU ≈ 12,000) and *E. coli* (Δ RFU ≈ 13,600) sustained significantly smaller viability drops, suggests a tighter regulation of free intracellular iron and antioxidant mechanism, which temporarily prevents the conversion of accumulated peroxides into fatal ^•^OH bursts highlighting endogenous redox buffering capacities.

### 3.3. Membrane permeabilization was linearly coupled to the extracellular release of siderophores

Partial reduction of molecular oxygen (O_2_) generates reactive oxygen species (ROS), including superoxide anions (O_2_^•−^), H_2_ O_2_, and hydroxyl radicals (^•^OH), which can promote lipid peroxidation and broader cellular injury ^34^. To determine whether oxidative membrane damage is associated with iron-acquisition capacity across distinct bacterial pathogens, we examined the relationship between inner membrane permeability, measured as log_10_ propidium iodide (PI) fluorescence in RFU, and siderophore production (%PSU) under baseline control and H_2_O_2_-challenged conditions (Figure 1d). Because PI is membrane-impermeant, increased PI fluorescence indicates dye entry into cells with compromised membranes. To quantify relationship between membrane damage and siderophore production, we modeled PI intake by cell (indicating membrane damage) as a predictor of extracellular siderophore release under baseline and peroxide-challenged conditions (Figure 1e). Under control conditions, simple linear regression demonstrated positive coupling between PI intake and resulting fluorescence and siderophore secretion (Control R^2^= 0.371). After exogenous H_2_O_2_ challenge membrane permeabilization (log_10_ [PI RFU]) remains tightly coupled to siderophore release, with membrane permeability explaining more than half of the variance in siderophore secretion (H_2_O_2_ R^2^ = 0.511).

### 3.4. Oxidative stress remodels iron, redox, and energy metabolism while reducing transport and redirecting evolution toward acid and peroxide defense in *E. coli*

To further understand the stress mechanism and elucidate the evolutionary trajectories of bacterial adaptation, genome-wide transcript abundances after H_2_O_2_ exposure in *E.coli* were tracked across baseline *E. coli* control. The RNASeq data obtained from Zorraquino et al., 2017 ^16^, was categorized into acute shock-unevolved (G0), and adaptively evolved *E. coli* after 500 generations (G500) exposed to oxidative stress. Differential gene expression analysis revealed widespread global transcriptomic restructuring, characterized by immediate acute stress responses followed by distinct metabolic re-routing in the adapted phenotype (Supplementary file S7-S9). Overall, transcript allocation among top differentially expressed genes (DEGs) shifted dramatically between conditions (Figure 1a). Ribosomal RNA genes (rrsB, rrfB) displayed massive up-regulation during acute shock that persisted into the adapted G500 state (*p_adj_* < 0.05). Conversely, primary stress-response genes like cspA were transiently down-regulated in both G0 and G500. Gene Ontology (GO) enrichment analysis confirmed distinct functional transitions across evolutionary phases(Figure 2b), the differential gene expression of G500 versus G0 demonstrated enrichment of central amino acid, tryptophan biosynthesis and iron-sulfur (Fe-S) cluster metabolic processes, including catalytic, lyase and transferase activity pathways indicating an adaptive shift from acute shock response to sustainable long-term homeostasis to protect against internal Fenton cytotoxicity.

**Figure 2.**
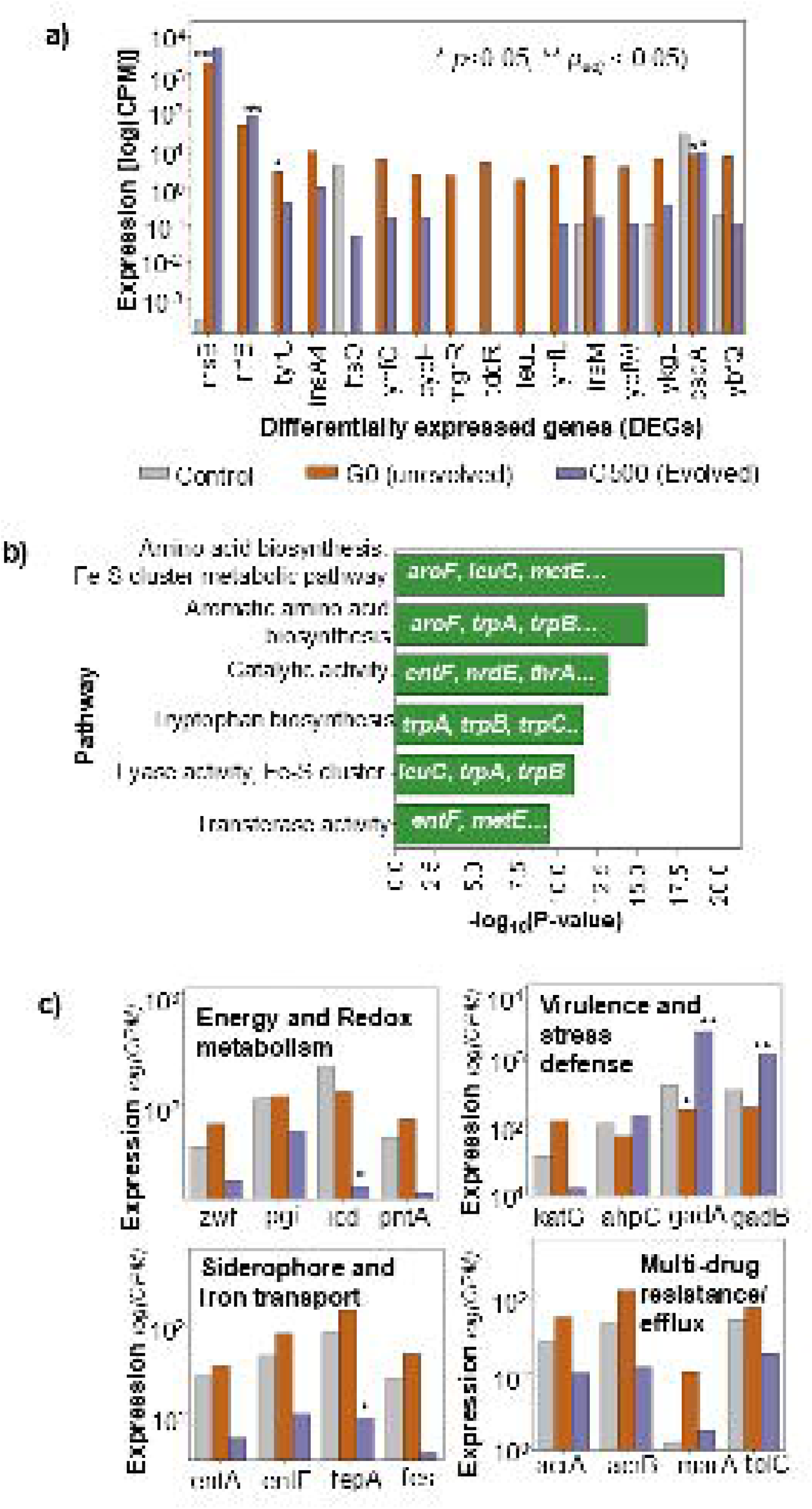
Transcriptomic reprogramming in unevolved acute shock (G0) vs. evolved long-term adapted (G500) E. coli states. **(a)** Gene expression profile log[Counts per million (CPM)] of top differentially expressed genes (DEGs) across control (gray), G0 acute shock (orange), and G500 adapted (purple) states (\**p* < 0.05, *p_adj_* < 0.05). **(b)** Gene Ontology (GO) functional enrichment mapping of DEGs highlights key metabolic, catalytic, and Fe-S cluster pathways (-log_10_[*p-value*]). **(c)** Specific transcriptomic shifts in energy/redox metabolism (zwf, pgi, icd, pntA), virulence/stress defense (katG, ahpC, gadA, gadB), siderophore/iron transport (entA, entE, fepA, fes), and multidrug efflux (acrA, acrB, marA, tolC).

Analysis of core energy and redox maintenance pathways revealed a clear transition from temporary stress survival to metabolic streamlining (Figure 2 c). Glucose-6-phosphate dehydrogenase (zwf) and pyridine nucleotide transhydrogenase (pntA) were elevated during acute oxidative shock (CPM increased from 40.1 to 66.4 for zwf, and 49.8 to 72.1 for pntA), supporting NADPH regeneration as observed in previous studies^35,36^ while in the adapted state (G500), it was down-regulated (pntA: 15.6 CPM; zwf: 20.3 CPM), indicating reduced reliance on acute redox-reducing equivalents. Further, TCA cycle optimization was observed by significantly downregulating isocitrate dehydrogenase (icd) from 221.3 CPM at control to 127.7 CPM in acute shock (G0), decreasing further to 10.0 CPM in adapted cells. Phosphoglucose isomaerase (pgi) maintained a stable baseline flux before downregulating in the evolved state (56.4 CPM versus 111.8 CPM control). Distinctly, the acute oxidative shock of G0 *E. coli* displayed non-significant up-regulation iron-acquisition and multi-drug efflux system (Figure 2c), while in G500 downregulation of major nonspecific porins, especially ompF (0.039-fold relative expression) and lamB (0.064-fold), with stable ompC expression was observed (Supplementary file S8-S9).

Likewise, following evolutionary adaptation to oxidative stress, siderophore machinery was strongly suppressed (fepA: 8.9 CPM, p_adj_ < 0.10; entE: 10.1 CPM; fes: 3.6 CPM). Similarly, multi-drug resistance/efflux system expression were reduced where, AcrAB-TolC efflux pump complex (acrA, acrB, tolC) and its global transcriptional activator marA were observed to be dynamically increased during acute shock (acrB CPM rose from 49.6 to 136.5; marA from 1.2 to 10.7). In adaptively evolved cells, expression across the entire complex returned to or fell below baseline levels (acrB: 12.3 CPM; acrA: 10.3 CPM; tolC: 18.7 CPM). It is noteworthy that while broad transport and central metabolic pathways were downregulated during adaptation (G500), specific enzymatic defense systems were heavily selected for long-term survival. On the other hand, an acid resistance system—Glutamate decarboxylases gadA and gadB demonstrated the most pronounced evolutionary shift. Suppressed during acute shock (gadA: 146.6 CPM; gadB: 160.1 CPM relative to 338.8 and 292.6 CPM control), both genes underwent massive transcriptional activation in adapted cells (gadA: 2237.7 CPM, LFC = 6.17, p_adj_ = 4.97 ×10^−9^; gadB: 1001.5 CPM, LFC = 4.87, p_adj_ = 1.15 ×10^−5^). Hydrogen peroxide scavenging enzymes displayed divergent specialization. Hydroperoxide reductase ahpC was up-regulated in adapted cells (120.1 CPM vs 89.6 CPM control), whereas catalase-peroxidase katG surged during acute shock (99.1 CPM) before being strongly suppressed in the evolved state (9.6 CPM).

### 3.5. Bifurcated transcriptional trajectories of iron acquisition and the fur–ryhB regulon under sustained ROS exposure

The accumulation of intracellular ROS driven by H_2_O_2_ in *E. coli* (obtained from, Figure 1 a (i)) was correlated with transcriptional activation in G0 across siderophore and other differentially expressed genes (Figure 3(a-b)). Within the siderophore pathway, gene expression partitioned into two distinct kinetic regimes: a highly responsive subset of 17 loci (R^2^ ≥ 0.50), including core iron acquisition and regulatory genes such as entB, entC, entD, fepA, fhuE, and fur, tracked intracellular ROS levels directly, shown in Figure 3a. Conversely, a second subset comprising 14 loci (R^2^ < 0.50) (Supplementary Figure S1), including entA, entE, fepB, fepC, fecA, and the small RNA ryhB, displayed expression profiles decoupled from ROS kinetics (rate of ROS accumulation inside cell). Cross-sample linear regression further demonstrated co-regulation between H_2_O_2_ detoxification and iron uptake machinery, evidenced by a significant positive correlation between the catalase-peroxidase katG and the ferric enterobactin outer membrane receptor fepA (R^2^=0.67, *p*=0.046, Figure 3c, Supplementary Figure S2 (a-c)). Additionally, on an individual locus level for G0 (acute oxidative stress), siderophore uptake and biosynthesis components (entF, fepA, cirA, fiu, fhuE, fur and rhyB) demonstrated robust positive Pearson correlations (*r*= 0.77-0.98) with primary oxidative stress response regulon members, including oxyR, katG, sodA, and gor, reinforcing an integrated transcriptional response during acute oxidative challenge (Figure3d).

**Figure 3.**
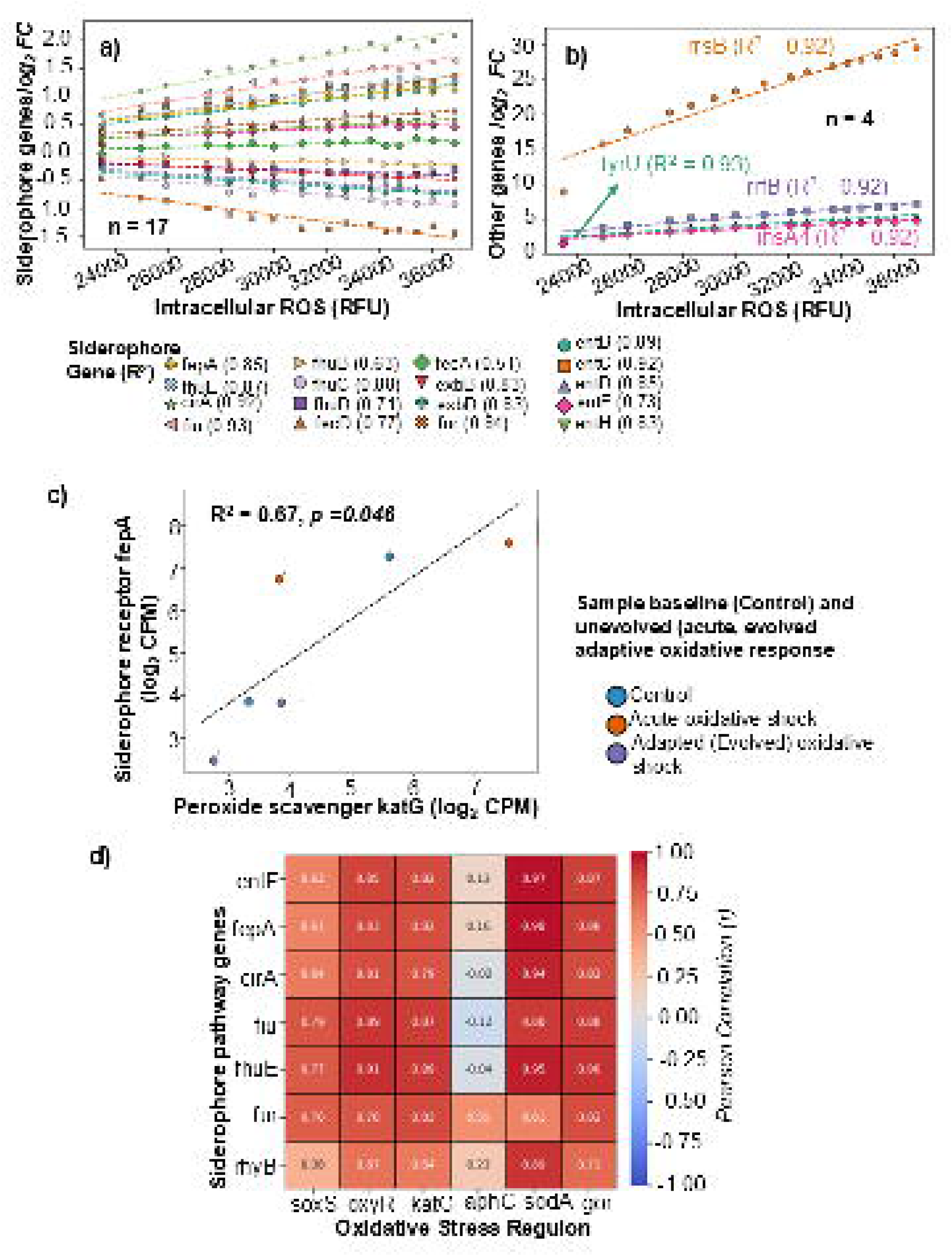
Co-expression kinetics and regulon coupling of iron acquisition pathways with oxidative stress response loci. **(a)**Linear correlation between intracellular ROS (RFU) and expression fold-change (log_2_ FC) for 17 siderophore-related genes. **(b)** Co-activation trajectories of non-siderophore marker loci (rrsB, tyrU, rrfB, insA4) relative to intracellular ROS levels (R^2^ ≥ 0.92). **(c)** Co-regulation regression plot of peroxide scavenger katG and siderophore receptor fepA. **(d)** Pearson correlation heatmap (r) evaluating co-regulation between individual siderophore loci (entF, fepA, cirA, fiu, fhuE, fur, rhyB) and primary members of the oxidative stress regulon (soxS, oxyR, katG, aphC, sodA, gor).

Despite acute co-activation, long-term evolutionary adaptation under sustained stress induced a pronounced transcriptional trajectory shift away from acute induction (Figure 4a). Siderophore receptor genes (fepA, fhuE, and cirA) and biosynthesis machinery (entF), which were activated during acute shock (log_2_FC ≈+0.5 to +2.1), underwent systematic suppression in the evolved state (log_2_FC≈ -1.8 to -3.0). Most notably, the iron-responsive small RNA regulator ryhB experienced an extreme downregulation, shifting from near-baseline levels during acute exposure (log_2_FC=-0.32) to marked repression in evolved lineages (log_2_FC=-8.28). Fisher’s exact, Chi-square, and hypergeometric over-representation analyses confirmed that siderophore pathway enrichment was specifically associated with the transition from acute stress to adaptive evolution (Evolved (G500) vs. Acute (G0)) rather than simple baseline comparisons against control conditions. Out of a total genome background of 4,440 loci containing 31 siderophore-related genes, 4 of the 23 DEGs identified in the evolved versus acute transition belonged to the siderophore pathway (Fisher’s Odds Ratio = 34.23, Fisher’s *p*=1.60×10^−5^; χ^2^=70.30, p=5.10× 10^−17^; Hypergeometric *p*=1.60×10^−5^)(Supplementary File S10). In contrast, no significant siderophore pathway enrichment was observed in acute vs. control (*p*=1.00) or evolved vs. control (p=0.387) comparisons (Supplementary File S10), demonstrating that down-regulation of iron acquisition machinery represents a highly specific evolutionary adaptation to attenuate Fenton-mediated oxidative damage. The selective down-regulation of iron import systems during long-term evolution reflects an adaptive trade-off: attenuating baseline iron uptake prevents excessive free intracellular Fe^2+,^ thereby mitigating Fenton-mediated •OH generation during sustained exposure to reactive oxygen species.

**Figure 4.**
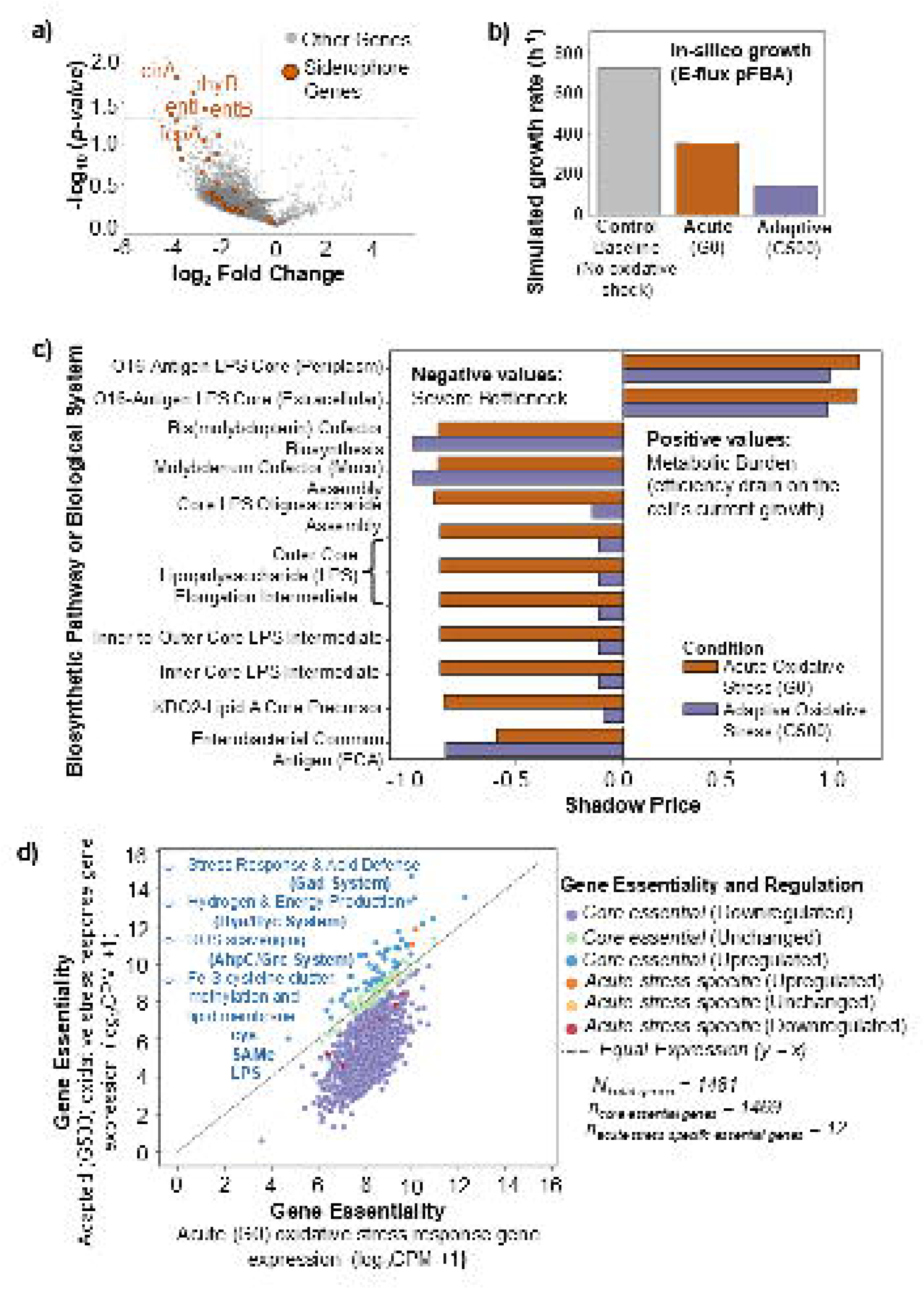
Metabolic flux balance, shadow price bottlenecks, and essentiality re-mapping during adaptive evolution. **(a)** Volcano plot showing global differential gene expression (log_2_FC vs. -log_10_[*p-value*]) highlighting the down-regulation of siderophore genes (orange circles) in adapted G500 populations. **(b)** In silico simulated growth rates (h^−1^) computed via E-flux parsimonious Flux Balance Analysis (pFBA) for control, acute (G0), and adapted (G500) states. **(c)** Shadow price analysis quantifying metabolic efficiency penalties (positive values) and severe pathway bottlenecks (negative values) across core LPS, cofactor, and cell envelope biosynthetic pathways. **(d)** Comparative gene essentiality plot (log_2_CPM + 1) comparing acute (G0) vs. adapted (G500) states, categorizing core essential (purple, green, blue) and acute-stress-specific essential genes (orange, yellow, red) (N_total_ = 1481).

### 3.6. Integration of E-Flux and constraint-based modeling identifies systems-level adaptations to long-term oxidative stress in *E. coli*

#### 3.6.1. Phenotypic growth trade-offs and capacity recovery under oxidative stress

To quantify the physiological impact of oxidative challenge and long-term evolutionary adaptation, genome-scale metabolic modeling of *Escherichia coli* (iML1515) was integrated with condition-specific transcriptomic fold-change profiles using the E-Flux algorithmic framework. Using the Parsimonious Flux Balance Analysis (pFBA) for baseline growth predictions, the model established an optimal unconstrained proliferation rate of 726.10 h^−1^ for the wild-type control. When subjected to acute oxidative shock (G0), expression constraints imposed by H_2_O_2_ exposure reduced the theoretical maximum growth rate by 50.9%, down to 356.46 h^−1^. This acute growth suppression reflects the immediate resource re-allocation toward cellular defense, enzyme repair, and ROS-scavenging pathways at the expense of central carbon assimilation. Following 500 generations of adaptive laboratory evolution under continuous oxidative selection, the G500 model demonstrated a further reduction in theoretical maximum biomass yield, reaching 144.39 h^−1^ (Figure 4b). Rather than restoring unconstrained baseline proliferation, the evolved regulatory network permanently prioritizes protective envelope maintenance, redox balance, and metabolic stability. This lower growth velocity represents an evolutionary trade-off where basal metabolic capacity is sacrificed to ensure structural resilience against persistent oxidative damage.

#### 3.6.2. Contraction of global flux flexibility and pathway streamlining

To assess how evolutionary adaptation reshapes the broader metabolic solution space, Flux Variability Analysis (FVA) performed across 2,712 reactions at a 95% suboptimal biomass threshold (f = 0.95). Comparing allowable flux spans (Δv_r_ = v_r,max_ -v_r,min_) between control, G0 and G500 revealed a systemic, genome-wide contraction in metabolic flexibility during long-term adaptation. While the acute shock state (G0) maintained broad flux ranges across diverse pathways—allowing the network to dynamically shift among alternative routes, the adapted state (G500) displayed a pronounced cluster of reactions shifting toward the origin (y < x, Supplementary Figure S3, Table 1).

**Table 1.**
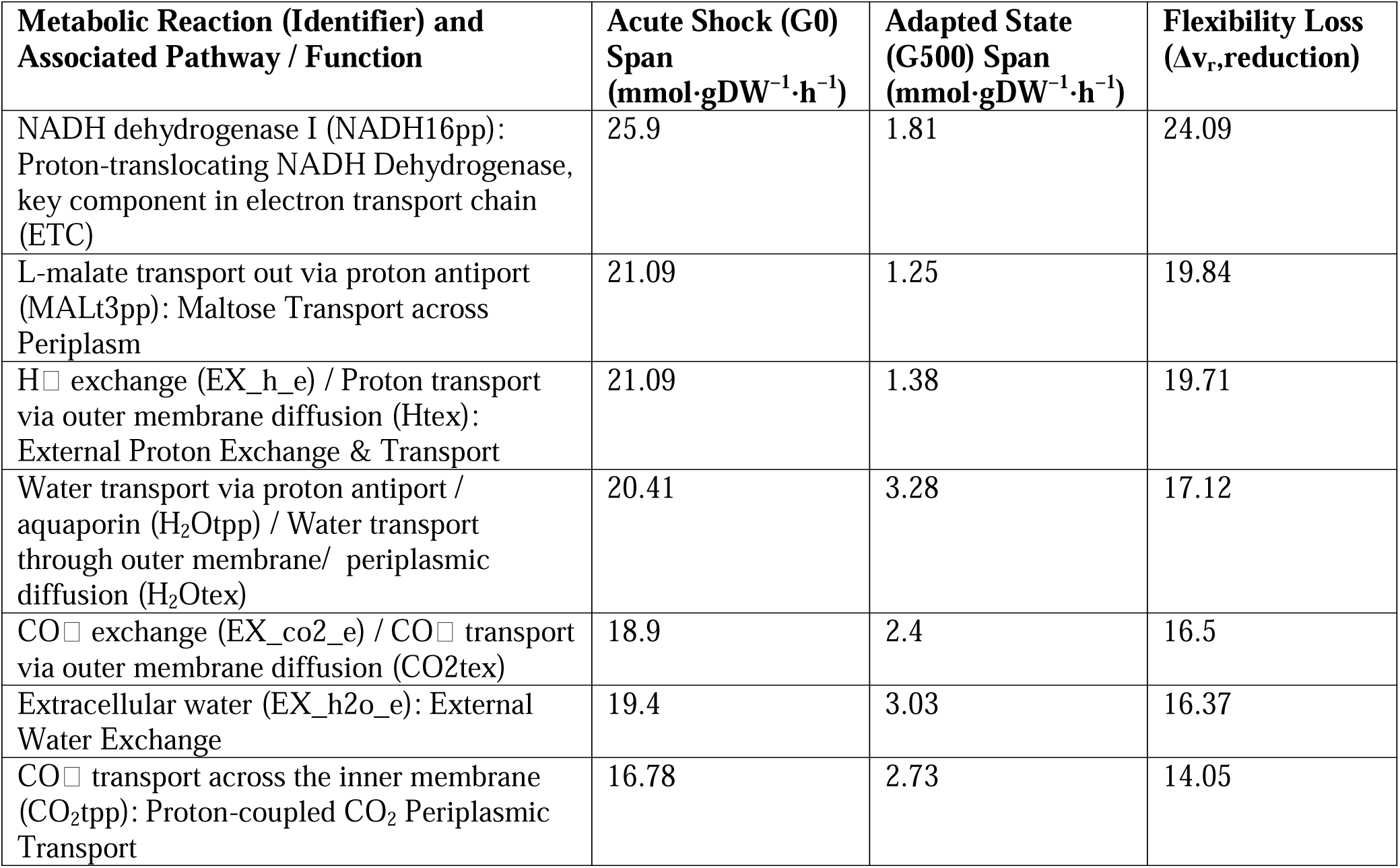
Genome-Scale Flux Variability Analysis (FVA) Contraction Across Acute Shock and Adapted Evolutionary States. Comparison of maximum permissible flux ranges mmol⋅ gDW^−1^⋅h^−1^ between acute oxidative shock (G0) and the long-term adapted state (G500) across primary metabolic pathways.

The largest loss of metabolic flexibility occurred within primary respiratory and proton-motive force (PMF) reactions. The NADH dehydrogenase I reaction (NADH16pp) exhibited the sharpest individual reduction, with its allowable flux span dropping from (25.90 mmol⋅gDW^−1^ h^−1^) during acute shock (G0) to just (1.81 mmol⋅gDW^−1^ h^−1^) in the adapted state (G500); (Δv_r_ = 24.09). A similar tightening was observed across external proton transport (EX_h_e, Htex; (Δv_r_ = 19.71) and respiratory byproduct exchange (EX_co2_e, CO2tex; (Δv_r_ = 16.50). This system-wide narrowing extended to dicarboxylic acid homeostasis via MALt3pp (Δv_r_ = 19.84), a key inner-membrane antiporter that regulates the PMF by exporting cytosolic L-malate in exchange for periplasmic protons. Collectively, this pathway streamlining demonstrates that long-term adaptation restricts metabolic operation into tightly regulated metabolic channels, minimizing redundant or energetically inefficient flux variations.

#### 3.6.3. Dual shadow price analysis and envelope bottleneck re-alignment

To elucidate the metabolic constraints and thermodynamic bottlenecks governing biomass accumulation across evolutionary states, a linear programming shadow price analysis was performed. Within this flux balance analysis (FBA) framework, shadow prices (λ) quantify the marginal sensitivity of the growth objective function to individual metabolite availability. Negative shadow prices identify growth-limiting precursors or substrates whose restricted availability directly bottlenecks biomass yield; conversely, positive shadow prices denote thermodynamic or structural demand constraints, where the accumulation and export of specific endpoint assemblies are strictly tied to cellular growth efficiency (Figure 4c, Supplementary file S13).Outer membrane lipopolysaccharide (LPS) assembly units—specifically periplasmic and extracellular O-antigen core complexes (o16a4colipa_p and o16a4colipa_e)—exhibited the highest positive shadow prices across both metabolic regimes, maintaining values of (λ = +1.102 and +1.093) during acute shock (G0) and settling at (λ = +0.965 and +0.955) in the adapted state (G500). This persistent positive demand confirms that the structural deployment of mature O16-LPS at the cell surface remains a strict, rate-limiting requirement for cellular replication. In the acute phase, biomass accumulation is heavily constrained by an operational bottleneck within the macromolecular Lpt translocation machinery. Upon long-term adaptation (G500), however, this export constraint was marginally alleviated, allowing metabolic control to be shared with downstream cellular systems.

Unlike to these structural export constraints, cytosolic inner-core lipopolysaccharide intermediates (colipa_c, gggagicolipa_c, gicolipa_c, ggagicolipa_c, gagicolipa_c, icolipa_c, and kphphhlipa_c) imposed severe metabolic penalties during immediate shock (λ ≈ -0.856 to - 0.884). This pronounced negative bottleneck underscores the steep energetic and precursor drain induced by urgent, Waa/Rfa operon-mediated envelope repair following initial membrane damage. Remarkably, in G500, these inner-core biosynthetic penalties relaxed substantially, dropping to λ ≈ -0.114 to -0.145. This structural reduction in shadow price magnitude demonstrates that evolved *E. coli* successfully streamlined its internal pathway throughput. By optimizing intermediate synthesis, the adapted strains alleviated acute membrane-associated bottlenecks and significantly minimized the metabolic cost of baseline envelope maintenance. In contrast, molybdenum cofactor derivatives (bwcogdp_c and bmocogdp_c) displayed increased negative shadow prices in the adapted state (λ=-0.986) compared to acute shock (λ=-0.865). While the lipopolysaccharide (LPS) core relaxes its metabolic burden over time, the metabolic synthesis cost of the molybdenum cofactor (*moa/mob/moe* genes) becomes significantly more expensive in the adapted state. Under long-term stress, the cell deliberately redirects a larger portion of its precursor pool toward Moco-dependent oxidoreductases. This increased drain contributes as a necessary investment to fuel systemic reactive oxygen species (ROS) neutralization and maintains internal redox balance. Similarly, the enterobacterial common antigen (ECA) precursor eca4und_p showed a heightened penalty in G500 (λ=-0.835) compared to G0 (λ=-0.589). In acute shock, the synthesis of the outer membrane glycans via the Wec operon is a secondary concern compared to the collapsing LPS core. However, once the LPS core assembly bottleneck is resolved at (G500), the ECA pathway emerges as a prominent metabolic drain. This increased penalty is driven by a critical resource conflict: both the expanding ECA network and the stabilized LPS pathways must compete for a limited pool of shared undecaprenyl phosphate (C55-P) lipid carriers, elevating the systemic cost of maintaining outer envelope stability.

#### 3.6.4. In Silico Gene Essentiality and Expression Remodeling Under Oxidative Stress Adaptation

Systematic in-silico single-gene deletion screens were performed to evaluate gene vulnerability and essentiality transitions across evolutionary stages. Knockouts yielding <10% of unmutated condition growth were classified as essential (Supplementary file S14). Out of a total N_total_ = 1481 analyzed genes, n_core_ essential = 1469 were classified as core essential genes, while n_acute_ stress specific = 12 were identified as acute-stress-specific essential genes. Comparative expression scatter analysis (log_2_(CPM + 1)) relative to the line of equal expression (y = x) revealed distinct regulatory trajectories among essential functional modules (Figure 4 d). Majority of the core essential genes (purple dots) clustered below the y = x parity line, demonstrating a widespread transcriptional downregulation in the long-term adapted state (G500) relative to acute exposure (G0). A distinct subset of essential genes was significantly upregulated (y > x) in the G500 adapted strain (blue dots), highlighting key mechanisms for oxidative stress tolerance.

i. The glutamate decarboxylase (Gad System) exhibited the highest upregulation, supporting pH homeostasis and secondary stress protection [*Stress Response and Acid Defense]*.
ii. The Hya/Hyc System (hydrogenases) was elevated, indicating a shift toward optimizing bioenergetic electron flux and redox balance [*Energy & Hydrogen Production]*.
iii. Peroxiredoxin/reductase complexes (AhpC/Grc System) were strongly induced to manage cellular peroxide burden continuously [*ROS Scavenging]*.
iv. Essential genes involved in iron-sulfur (Fe-S) cysteine cluster assembly, S-adenosylmethionine (SAMe) methylation pathways, and lipopolysaccharide (LPS) cell envelope integrity (cys, SAMe, LPS) showed sustained induction to repair damaged cellular machinery [*Membrane and Redox Cofactor Biosynthesis]*.
v. Acute-stress-specific essential genes displayed variable expression patterns (red, orange, and light-yellow dots), with specific candidates downregulated in G500, pointing to a transition from acute survival mechanisms to a streamlined, adapted metabolic state [*Acute Stress-Specific Genes]*.

## Discussion

Our study demonstrated that bacterial survival under H_2_O_2_ stress requires coordinated radical scavenging, structural repair, and regulation of intracellular labile iron pools (Figure 5). Our findings support a sequence in which peroxide exposure increases intracellular H_2_O_2_; excess labile Fe^2+^ converts it into hydroxyl radicals (^•^OH) through Fenton chemistry; and the resulting ROS burden correlates with membrane damage, viability loss, and transcriptome-supported repression of iron acquisition in *E. coli*. Thus, although siderophores are essential high-affinity iron scavengers during nutrient limitation, active iron acquisition under peroxide challenge becomes a pro-oxidant liability by expanding intracellular iron pools, intensifying Fenton chemistry, and increasing oxidation surges. Across *A. baumannii*, *E. coli*, *P. aeruginosa*, and *S. aureus*, acute peroxide shock significantly increased intracellular oxidation (*p* < 0.0001; Figure 1a) and was accompanied by species-dependent suppression of growth-normalized extracellular siderophore output (Figure 1b). These differences suggest that baseline iron-acquisition activity and intrinsic redox buffering jointly shaped oxidative outcomes: high baseline siderophore producers such as *P. aeruginosa* showed elevated internal cellular oxidation, whereas stronger redox-buffering species such as *S. aureus* more effectively restricted cellular oxidation surge during peroxide challenge (Figure 1c).

**Figure 5.**
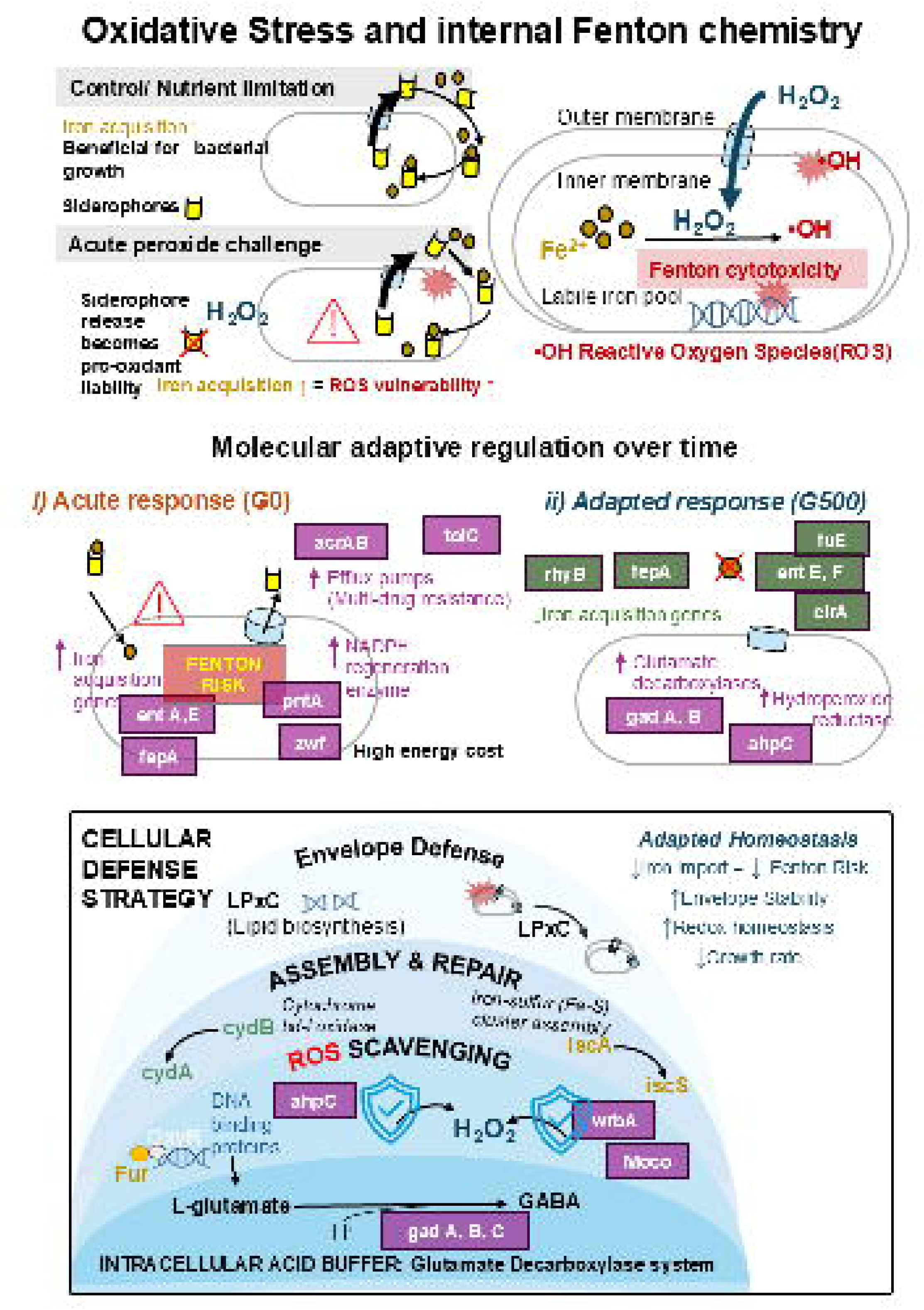
Conceptual model of internal Fenton toxicity and systemic cellular defense strategies during long-term adaptation. (Top) Mechanistic overview of Fenton chemistry: siderophore-driven iron acquisition enhances growth in nutrient-limited environments but becomes a lethal pro-oxidant liability during acute peroxide challenge. (Middle) Timeline of molecular adaptive regulation, contrasting high-energy-cost acute responses (G0; upregulation of entA/E, fepA, pntA, zwf, and acrAB-tolC) with optimized long-term adaptation (G500; suppression of iron acquisition alongside induction of gadA/B and ahpC). (Bottom) Integrated defense architecture in adapted homeostasis (G500), coordinating envelope stabilization (lpxC), ROS scavenging (ahpC, wrbA), Fe-S cluster/protein repair (iscA/S, cydA/B), and acid buffering via the glutamate decarboxylase system (gadA/B/C).

Our study established, cellular injury evidence through real-time uptake of Propidium Iodide (PI). PI fluorophore exclusion depends directly on membrane structural integrity; its progressive intracellular entry provides physical proof of lipid peroxidation and envelope permeabilization induced by hydroxyl radical generation during Fenton turnover. Further at the molecular level, persistent transcriptional induction across core stress regulons served as an unbiased biological reporter of intracellular oxidative burden. Differential expression analysis revealed sustained upregulation of primary peroxide scavengers (*ahpC*), acid/glutamate-dependent homeostasis systems (*gadA, gadB*), and iron-sulfur cluster repair machinery (*iscA, iscS*). These transcriptional signatures confirm active physiological defense and enzymatic damage repair independent of exogenous chemical indicator kinetics. To evaluate the systemic cost of oxidative stress response, E-Flux-integrated Flux Variability Analysis (FVA) demonstrated a global contraction in metabolic flexibility. Shadow price optimizations identified severe energetic bottlenecks and metabolic re-routing enforced by the cell’s requirement to maintain reducing equivalents (NADPH) and repair damaged cofactors. This constraint-based analysis confirms that oxidative defense imposes a measurable, system-wide metabolic trade-off that dictates survival under hydrogen peroxide exposure.

### Physiological Trade-Off Between Iron Acquisition and Oxidative Vulnerability

Acute H_2_O_2_ stress triggers Fur/OxyR-mediated siderophore downregulation to limit Fenton-driven •OH generation, but species with strong antioxidant defenses, such as *P. aeruginosa*, can sustain iron acquisition more safely than species with lower antioxidant capacity ^36,37^. By integrating moderation regression with viability (log_10_ CFU/mL) and membrane-integrity metrics, we identified a conditional trade-off: H_2_O_2_-exposed bacteria reduced siderophore output (Figure 1b), whereas species with higher baseline siderophore production survived better during acute oxidative stress despite net intracellular ROS increases (Figure 1c). In *S. aureus*, oxidation amplification corresponded with reduced viable colony counts, while *P. aeruginosa* likely buffered Fenton-derived ROS through robust antioxidant systems, retaining siderophore release without equivalent viability loss. Thus, siderophore production may act as an evolutionary double-edged sword, supporting competitive iron acquisition when oxidation is controlled but increasing oxidative vulnerability when peroxide and labile iron converge.

The link between iron uptake, radical generation, and membrane damage was further supported by strong linear coupling between inner membrane permeability (log_10_ PI [RFU]) and extracellular siderophore capacity (Figure 1e). This pattern suggests that peroxide damages membrane lipids and weakens envelope barrier function, allowing passive chelator leakage and altering extracellular siderophore detection, while continued iron-siderophore uptake can increase intracellular Fe^2+^ and amplify Fenton-mediated injury. Membrane permeability consistently predicted siderophore release across fixed-effects and linear mixed-effects models, indicating that envelope destabilization closely associates with extracellular siderophore release. In *E. coli*, surge in intracellular oxidation and oxidative-stress regulons positively correlated with siderophore genes (Figures 3 (a-c), 4a), and adaptive downregulation and enrichment of siderophore biosynthesis limited iron import and Fenton-mediated damage (Supplementary File S9 and S10).

Iron-siderophore imports through TonB-dependent transporters (*fepA, fhuE, cirA*) and ABC transporters (fepCDB) can fuel intracellular Fenton chemistry ^38–40^; therefore, repressing siderophore biosynthesis and iron import (*entF, entE, fes*) directly lowers •OH production. Species with the highest relative ROS spikes, such as *E. coli*, rapidly reduced extracellular chelator release. To define the molecular basis of oxidative stress response, we integrated experimental phenotypes, transcriptomics, and genome-scale metabolic modeling (iML1515) across acute unevolved shock (G0) and 500-generation adapted *E. coli* lineages (G500). These approaches revealed a stepwise adaptive trajectory: acute peroxide exposure activated emergency defenses, including NADPH-regenerating enzymes (*zwf, pntA*), multidrug efflux systems (*acrAB-tolC, marA*), and iron-transport/siderophore machinery (*fepA, entA, entE*), which may help repair oxidized iron-sulfur (Fe-S) clusters but can also expand labile iron and intensify Fenton chemistry. Over evolution, this costly emergency state shifted toward energy conservation and iron suppression. In G500, siderophore receptor loci (*fepA, fhuE, cirA*) and biosynthesis genes (*entF, entE, fes*) were strongly repressed, while *ryhB* showed marked transcriptional collapse (Log_2_FC = -8.28). Enrichment testing confirmed that siderophore-pathway downregulation was specific to the adaptive transition from acute shock to evolution (Evolved versus Acute; Fisher’s Odds Ratio = 34.23, *p* = 1.60 ×10^−5^). These results support a model in which sustained oxidative selection favors reduced iron acquisition because limiting labile Fe^2+^ lowers Fenton-mediated hydroxyl-radical production and improves long-term cellular stability.

Because evolved cells suppress iron acquisition and high-cost transport, they redirect resources toward redox balance and envelope integrity. Evolved *E. coli* strongly upregulated acid-resistance glutamate decarboxylases (gadA, gadB; Log_2_FC = +6.17 and +4.87) and hydroperoxide reductase *ahpC*, while reducing reliance on energy-intensive efflux pumps and transient NADPH overproduction. pFBA modeling showed that this protective reallocation reduced theoretical maximum growth from 726.10 h^−1^ in unstressed wild type to 356.46 h^−1^ during G0 shock and 144.39 h^−1^ in G500, indicating a shift from rapid proliferation toward survival. FVA across 2,712 reactions further showed narrowed metabolic flexibility, especially in proton-translocating NADH dehydrogenase (NADH16pp; Δv_r_ loss = 24.09 mmol gDW^−1^h^−1^) and proton exchange reactions, channeling metabolism into tightly regulated, efficient routes. Shadow price analysis showed that this re-routing relieved acute membrane bottlenecks, decreasing cytosolic inner-core lipopolysaccharide precursor penalties from λ ≈ -0.87 in G0 to λ ≈ -0.13 in G500.

In-silico single-gene essentiality mapping showed that evolved *E. coli* eliminated all 12 acute-specific conditional vulnerabilities, including core glycolytic enzymes (gapA, eno) and ATP synthase subunits (atpA-H). Thus, over 500 generations, *E. coli* reduced Fenton risk, re-routed metabolic flux, relieved envelope bottlenecks, and established bypass routes that improved genetic robustness under continuous oxidative stress. During acute shock, *E. coli* induced resource-intensive siderophore systems, including b0180 and b0881, to acquire iron for Fe-S cluster repair; in the evolved state, enterobactin and transport genes, including *entF* and f*epA*, were downregulated relative to acute levels. This reduction limits intracellular iron accumulation and Fenton-mediated damage while lowering the energetic cost of iron-transport overproduction. Long-term adaptation therefore replaces broad transcriptional overactivation with targeted maintenance of protective systems, including *asr, gadA, dps,* and *dnaK*, establishing an energy-efficient survival baseline.

### Re-routing of core essential genes and redox homeostasis in the adapted survival state

The shift from acute exponential growth to a resilient, stationary-like adapted state reflects coordinated rewiring of core pathways. Rather than entering nonspecific metabolic arrest, cells reorganize transcription to strengthen the envelope, neutralize oxidative stress, suppress Fenton chemistry, and limit antibiotic entry. A key feature is activation of the glutamate decarboxylase system, including *gadA* (26.12-fold), *gadC* (11.10-fold), and *gadB* (10.67-fold), which consumes intracellular protons during conversion of L-glutamate to GABA and exports GABA through GadC, thereby buffering cytoplasmic acidification, preserving membrane potential, and supporting survival. The adapted transcriptome also induces layered ROS and electrophile defenses: *ahpC* (5.41-fold) scavenges endogenous H_2_O_2_, while wrbA (6.50-fold) promotes two-electron quinone reduction, limiting toxic semiquinone intermediates and downstream superoxide generation. Activation of cytochrome *bd*-I oxidase subunits *cydA/cydB* and Fe-S cluster assembly genes *iscA/iscS* further preserves respiratory electron flow and repairs oxidized Fe-S proteins during microaerobic or stagnant growth.

This metabolic reorganization also promotes nonspecific multidrug cross-resistance through outer membrane remodeling. Downregulation of major nonspecific porins, especially *ompF* (0.039-fold relative expression) and *lamB* (0.064-fold), with stable *ompC* expression, reduces overall permeability and restricts entry of hydrophilic antibiotics such as β-lactams and fluoroquinolones. Upregulation of *lpxC* further supports lipopolysaccharide biosynthesis and envelope reinforcement, allowing passive antimicrobial resistance without classical acrA, acrB, or *tolC* efflux overexpression. Entry into the adapted state also requires strict limitation of intracellular Fenton chemistry: Fe^2+^ reacts with H_2_O_2_ to generate ^•^OH, causing lipid peroxidation, DNA strand breaks, and protein carbonylation. To prevent this cascade, adapted cells suppress iron acquisition, including *entA, fepA, fhuA,* and *bfr*, while tightly regulating *feoB*. By limiting labile iron influx and stabilizing remaining iron pools, adapted bacteria decouple iron physiology from metabolic ROS production. This Fur-, OxyR-, and RpoS-shaped response supports long-term persistence while maintaining viability and structural integrity under prolonged oxidative stress.

## 4. Conclusion

This study demonstrates that bacterial adaptation to oxidative stress is governed by a tightly coordinated axis linking iron homeostasis, envelope integrity, and central metabolic flux. Across diverse bacterial pathogens, acute hydrogen peroxide exposure triggers an immediate physiological shift, dampening growth-normalized siderophore production to prevent membrane permeabilization and cellular oxidation amplification. Oxidative membrane damage acts as the primary driver of passive siderophore leakage, which cells counter by suppressing siderophore biosynthesis to restrict iron influx. In *E. coli*, long-term evolutionary adaptation under sustained peroxide selection permanently reinforces this mechanism through the transcriptomic repression of siderophore biosynthesis and transport systems (*ent, fep, ryhB*), driving 56.5% of responsive sub-network loci back to a homeostatic baseline. Constraint-based metabolic modeling reveals that this iron-suppressive adaptation incurs a deliberate fitness trade-off: evolved strains restrict global flux variability and reduce maximal proliferation rates to resolve envelope bottlenecks, eliminate conditional single-gene essentialities, and preserve long-term redox balance. Ultimately, coordinated siderophore suppression and redox-metabolic realignment represent a fundamental bacterial survival strategy that balances nutrient acquisition against Fenton-mediated cytotoxicity during oxidative challenges.

## Supporting information

Suptary Figire

## CRediT authorship contribution statement

**Elise Bulaoro**: Data Curation, Methodology, Formal Analysis; **Bharat Mishra**: Data Curation, Methodology, Writing: review & editing; **Anuradha Goswami**: Fund acquisition, Conceptualization, Supervision, Methodology, Writing-original draft, review & editing.

## Acknowledgements

This study has received University of Notre Dame Internal funding support. We thank Dr. Michael J. McConnell at the University of Notre Dame for generously providing the *Acinetobacter baumannii ATCC 19606, Escherichia coli MG1655, Pseudomonas aeruginosa PAO1, and Staphylococcus aureus USA300* and laboratory resources used in this study.

## Supplementary Data

Supplementary data files and supplementary figures are available in Supplementary documents.

## Data availability

Additional Data and codes used to generate plots will be made available on request.

## Ethics declaration

The author(s) declares that the study does not involve humans nor animal subjects. The author(s) used Google Gemini to improvise the codes used in this study.

