## Supplementary material for "Adaptive siderophore repression and redox-metabolic remodeling limit *E. coli* Fenton-mediated cytotoxicity after H_2_O_2_ exposure": Suptary Figire

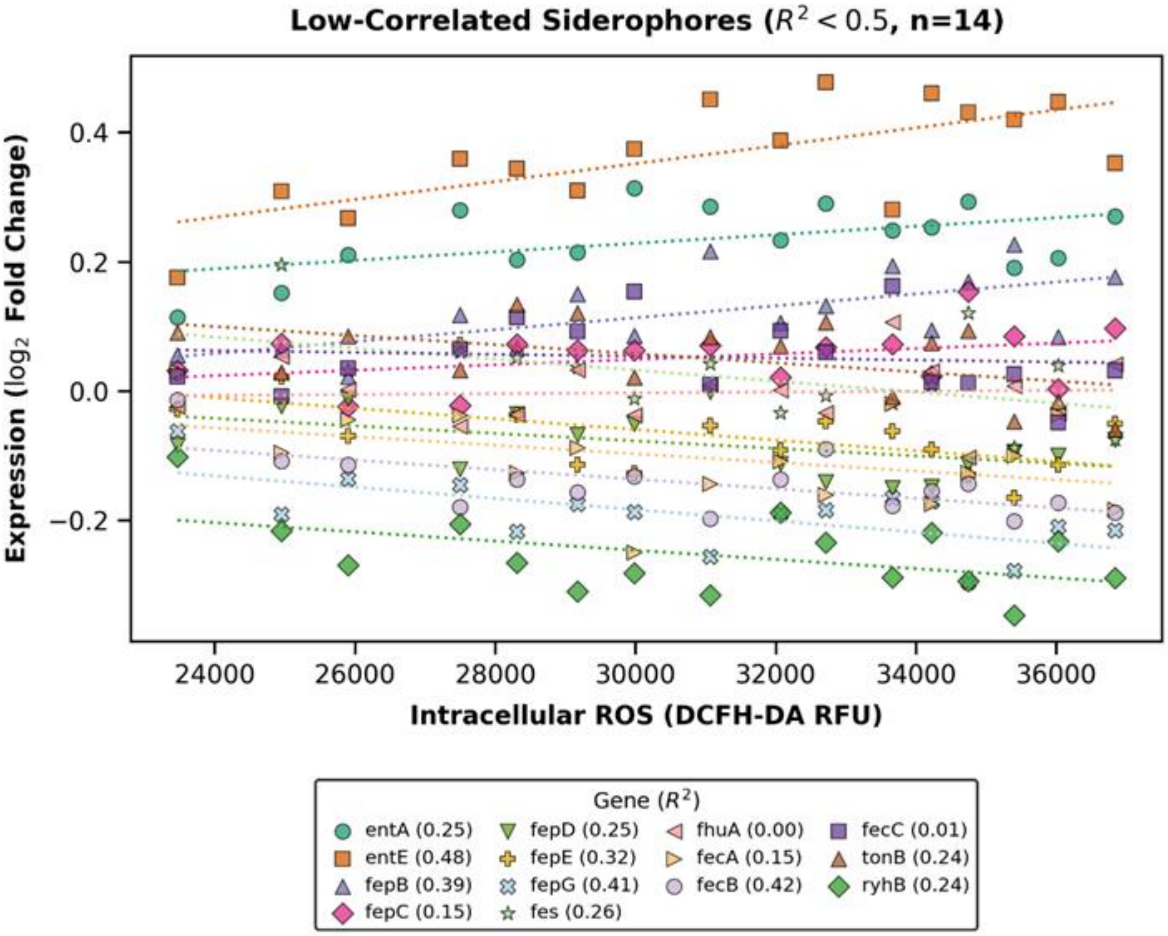


**Figure S1** Low-correlated siderophore genes in Acute Shock *E.coli* with intracellular Reactive Oxygen Species Generated during acute peroxide shock.

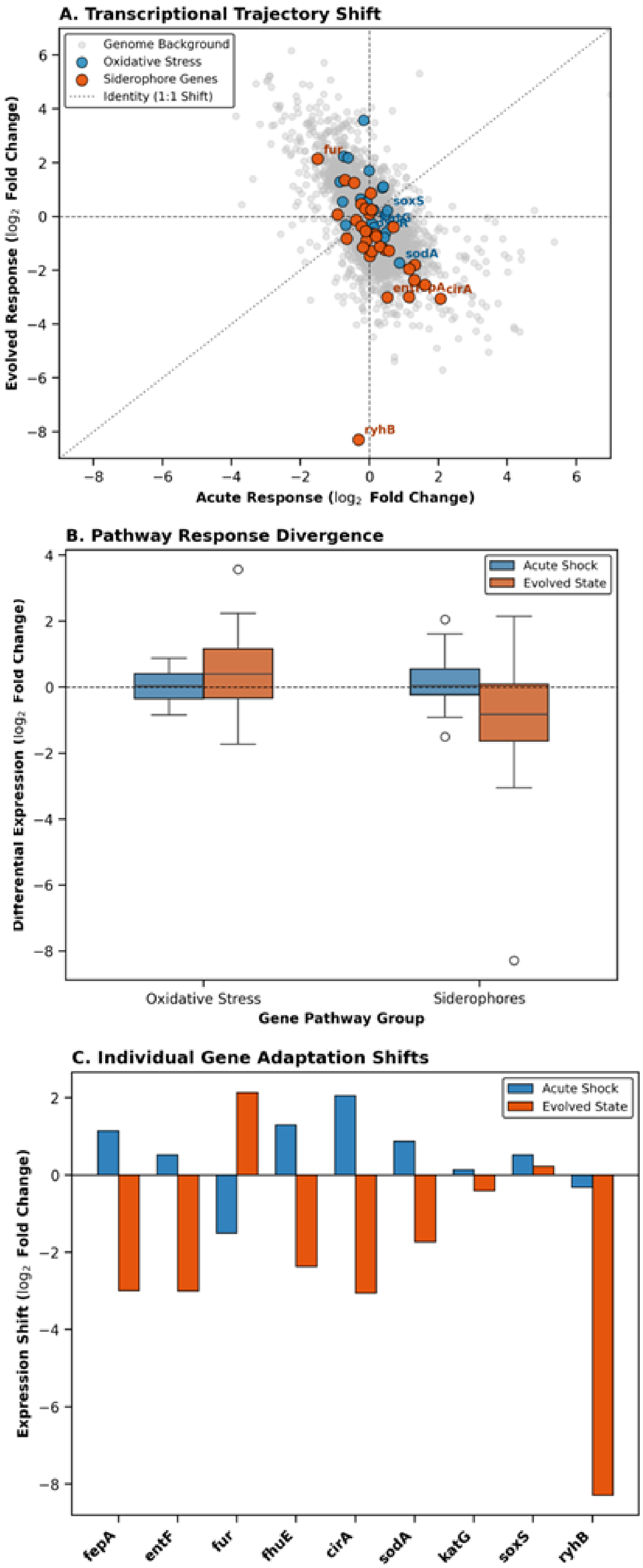


**Figure S2.** Divergent Transcriptional Reprogramming of Oxidative Stress Response and Siderophore Systems During Evolutionary Adaptation.

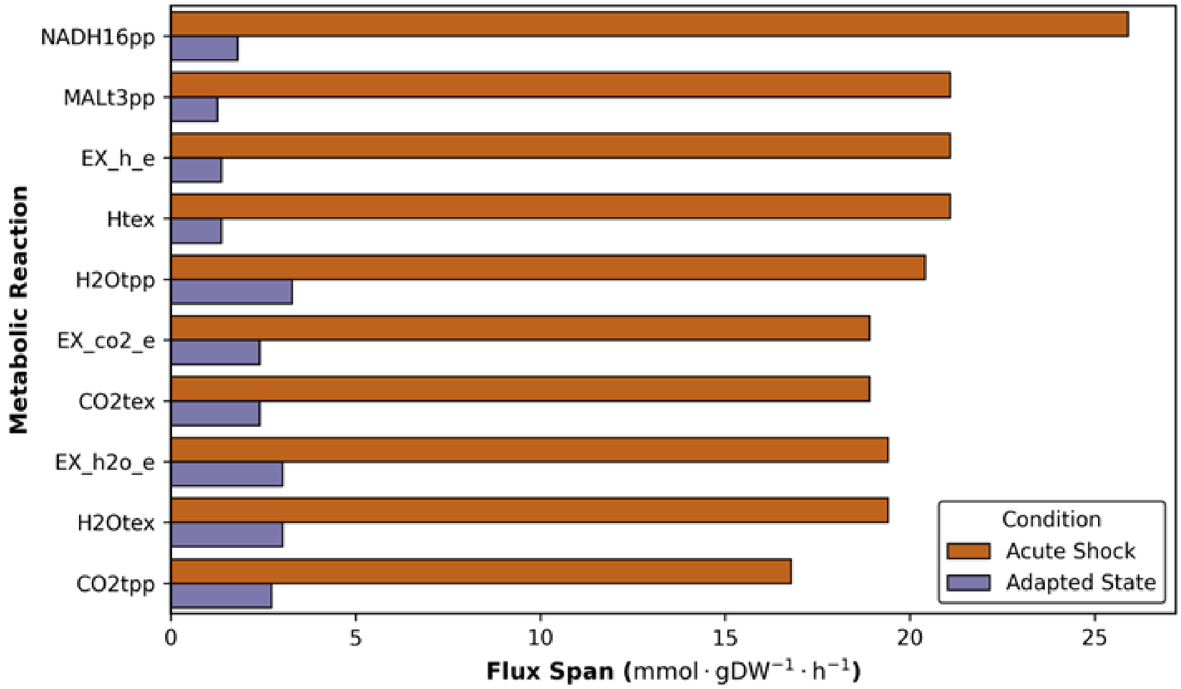


**Figure S3** Top streamlined reaction with largest loss of Flux flexibility
